# Neural correlates of successful relational memory in younger and older adults: An activation likelihood estimation meta-analysis

**DOI:** 10.64898/2026.05.18.725649

**Authors:** Min Sung Seo, Nancy A. Dennis

**Affiliations:** Department of Psychology, The Pennsylvania State University, University Park

**Keywords:** fMRI, cognitive aging, meta-analysis, associative memory, source memory, relational memory, activation likelihood estimation

## Abstract

Relational memory, the ability to encode and retrieve associations among multiple elements of an experience, is a core component of episodic memory that shows disproportionate age-related decline. Despite a substantial neuroimaging literature examining relational memory in aging, findings remain heterogeneous, and no quantitative synthesis has been conducted. The present study addressed this gap using Activation Likelihood Estimation (ALE) meta-analyses to characterize the neural correlates of relational memory success during encoding and retrieval in younger and older adults. Separate within-age-group, conjunction, and subtraction analyses were conducted, along with an exploratory analysis examining a general relational memory network across 70 independent studies. During encoding, younger adults showed robust convergence across medial temporal and prefrontal regions, whereas older adults showed more limited convergence. Shared convergence across age groups was observed in the left hippocampus and right inferior temporal gyrus, and direct age-group contrasts revealed greater prefrontal convergence in younger relative to older adults. During retrieval, younger adults showed convergence in posterior default mode and subcortical regions, whereas older adults showed convergence in the left angular gyrus, with no shared convergence observed across age groups. Across all studies, the hippocampus showed the most robust bilateral convergence across age groups and memory phases, underscoring its critical role in relational binding. Together, these findings provide the first quantitative characterization of the neural correlates of relational memory success in aging and highlight stable hippocampal involvement alongside age-related variability in prefrontal and posterior retrieval-related recruitment.

## 1 Introduction

Relational memory refers to the ability to encode and retrieve associations among multiple elements of an experience, including relationships between items (e.g., word–word or object–object associations) and links between items and their contextual features (e.g., spatial location, temporal context, or perceptual details, Davachi, 2006; Giovanello & Dew, 2015). Relational memory is a core component of episodic memory, the memory system that supports recollection of specific past events, including what happened, where it occurred, and when it took place (Tulving, 2002). Although memory for individual items provides a foundational building block of episodic memory, the relational component is essential for the richness of episodic memory. As such, episodic memory relies on the successful encoding and retrieval of relational information, defined here as memory for associations among items and between items and their contextual features.

Previous research has investigated relational memory primarily through two domains: associative memory and source memory. Associative memory refers to memory for relationships between items, such as remembering that two words or a face-name pair were previously studied together (De Brigard et al., 2020; Naveh-Benjamin et al., 2004, 2009). In contrast, source memory refers to memory for contextual features of an item, such as the location or modality in which an item was encountered (Glisky et al., 2001; Schacter et al., 1991; Uncapher et al., 2006).

Although these domains are often studied separately, they both rely on relational binding processes that link multiple elements into a unified memory representation and are supported by overlapping medial temporal and frontoparietal control networks. Thus, associative and source memory can be conceptualized as different expressions of a broader relational memory system.

Relational memory is more demanding than memory for individual items, even in younger adults, because it requires the encoding of multiple elements as well as the associations that bind them together (Craik & Byrd, 1982; Naveh-Benjamin, 2000; Old & Naveh-Benjamin, 2008). These additional representational and control demands often require recollection-based processes to successfully retrieve bound information (Diana et al., 2007; Prull et al., 2006; Yonelinas, 2002). Consistent with these demands, healthy aging is associated with disproportionate impairments in relational memory relative to memory for individual items (Chalfonte & Johnson, 1996; Naveh-Benjamin, 2000; Old & Naveh-Benjamin, 2008; Spencer & Raz, 1995; for reviews, see Dennis & McCormick-Huhn, 2018; Giovanello & Dew, 2015; Sümer & Kaynak, 2025). Although relational memory is generally more difficult than single-item memory across both younger and older adults, older adults show disproportionately greater deficits despite relatively preserved item memory (Naveh-Benjamin, 2000; Old & Naveh-Benjamin, 2008).

These relational deficits are observed across a wide range of associative paradigms, including word–word (Addis et al., 2014; Giovanello et al., 2012; Kim & Giovanello, 2011a; Wang & Giovanello, 2016), word–face (Steinkrauss et al., 2023), face–name (James et al., 2008; Naveh-Benjamin et al., 2004, 2009), face-scene (Chen & Naveh-Benjamin, 2012; Dennis et al., 2014), and item–item (Dennis et al., 2025).

Similarly, aging disproportionately impairs source memory relative to item memory (Chalfonte & Johnson, 1996; Glisky & Kong, 2008; Kuhlmann & Boywitt, 2016) across a variety of item-source pairings, including word–source (e.g., color, location, or voice; Foster et al., 2016; Swick et al., 2006), face–source (e.g., occupation, scene, or name; Dennis et al., 2008; Geier et al., 2020; James et al., 2019), and item–source (e.g., location; Siegel & Castel, 2018). Critically, across these paradigms, memory deficits are consistently localized to the relational component of memory, whereas item memory often remains relatively preserved. Together, these findings suggest that relational memory is particularly vulnerable to aging because successful performance depends on binding multiple elements into integrated representations.

### 1.1 Neural correlates of relational memory in aging

Despite clear behavioral evidence that relational memory is particularly demanding and disproportionately affected by aging, its neural correlates are far less consistent. Functional magnetic resonance imaging (fMRI) studies have examined relational memory across both encoding and retrieval, allowing researchers to assess whether variability arises during the formation of relational representations, their subsequent retrieval, or both. However, findings across studies and phases are not uniform, making it difficult to identify a consistent set of neural regions supporting relational memory in aging.

#### 1.1.1 Encoding

During relational encoding, successful memory formation is typically supported by a distributed network including the hippocampus, which supports relational binding, as well as prefrontal and posterior parietal control regions that contribute to strategic encoding and representational processing (Davachi, 2006; Ritchey et al., 2015; Yonelinas et al., 2019). Despite broad consensus regarding the involvement of these regions, findings across individual studies remain variable, likely reflecting differences in the cognitive processes emphasized across relational encoding paradigms. For example, some tasks place greater demands on semantic elaboration or organizational processing, often engaging the left ventrolateral prefrontal cortex, whereas others rely more heavily on perceptual associative binding or contextual integration, potentially emphasizing posterior medial temporal structures (Kim, 2023, 2024a). Variability in stimulus domain (e.g., verbal vs. visuospatial), encoding strategy (e.g., intentional vs. incidental encoding instructions), and the specific memory contrasts used to isolate successful encoding (e.g., comparing source-correct trials with item-only vs. forgotten trials) may further contribute to these inconsistencies. In addition, some paradigms differ in the extent to which they promote unitization (i.e., encoding multiple elements as a single coherent representation), a process thought to rely more heavily on the perirhinal cortex and potentially reduce reliance on hippocampal relational binding mechanisms (Diana et al., 2007; Haskins et al., 2008). Consequently, it remains unclear which neural regions show the most reliable and consistent convergence during successful relational encoding in younger adults.

In aging, these patterns become further complicated by age-related alterations in neural recruitment, with studies reporting both increased and decreased neural recruitment in older relative to younger adults. On the one hand, several studies report over-recruitment of neural activity in older adults during relational memory tasks (de Chastelaine et al., 2011; Giovanello & Dew, 2015; Miller et al., 2008). For example, in associative memory paradigms, younger adults show stronger task-related deactivation in default mode regions (e.g., medial parietal lobes) whereas older adults exhibit reduced or absent deactivation (de Chastelaine et al., 2011; Miller et al., 2008). In addition, older adults show increased recruitment of frontal regions and more bilateral activation patterns, which in some cases are associated with better associative memory performance (Bangen et al., 2012; de Chastelaine et al., 2016).

On the other hand, other studies report age-related under-recruitment of neural regions supporting relational encoding, including the hippocampus and prefrontal cortex. For example, Dennis and colleagues (2008) found age-related reductions in hippocampal and prefrontal activity during successful encoding of face–scene associations, and Kim & Giovanello (2011b) reported reduced activity in prefrontal, inferior parietal, and left hippocampal regions in older adults during successful encoding of word pairs. Such under-recruitment has been interpreted as reflecting reduced efficiency in hippocampal-prefrontal processes supporting relational binding and strategic encoding processes necessary for the formation of relational memories.

Importantly, age-related differences are not uniformly characterized by simple increases or decreases in neural recruitment. For example, some source memory encoding studies show age-related preservation of hippocampal engagement alongside age-related reductions in the recruitment of frontal control and posterior perceptual regions, coupled with increased engagement of other regions (e.g., occipital cortex or default mode network). This complex pattern has been interpreted as reflecting less efficient or less selective processing in older adults (Cansino, Estrada-Manilla, et al., 2015; Dulas & Duarte, 2011; Foster et al., 2016; Park et al., 2013).

These discrepancies have often been attributed to age differences in task demands and the type of relational information emphasized across paradigms. For example, Dennis et al. (2019) showed that age-related differences in associative encoding vary as a function of the type of association being processed (e.g., item–item vs. item–context), with distinct patterns of neural recruitment observed across sensory and medial temporal regions. Similarly, when relational encoding is supported by strategic processes such as self-referential encoding, older adults can recruit neural regions comparable to those engaged by younger adults (Leshikar & Duarte, 2014).

Under increased task demands, however, older adults often exhibit greater recruitment of frontoparietal control regions alongside reduced modulation of activity as cognitive demands increase (Kennedy et al., 2015; Spaniol & Grady, 2012). In addition, stimulus familiarity can alter relational encoding processes, with prior familiarization reducing activity across hippocampal, parahippocampal, and prefrontal regions while still supporting successful associative memory formation (Dennis et al., 2015; Elbich et al., 2021; Kremers et al., 2014; Vannini et al., 2013).

These findings can also be interpreted within relational binding frameworks, which propose that the hippocampus supports the integration of item and contextual information into unified representations through interactions with perirhinal and parahippocampal cortices (Davachi, 2006; Olsen et al., 2012; Ranganath, 2010; Yonelinas et al., 2019). From this perspective, age-related differences in neural recruitment may reflect differences in the quality of item and contextual representations available for binding, as well as differences in strategic and control processes that support integration. Thus, variability across studies may not necessarily reflect contradictory findings, but differences in the cognitive operations emphasized across relational encoding paradigms. Despite these proposed explanations, however, no consistent pattern of neural convergence during relational encoding has emerged across studies. As a result, it remains unclear which neural regions are reliably associated with successful relational encoding within each age group, which regions are shared across age groups, and where age-related variability in neural recruitment most consistently emerges across tasks and experimental contexts.

#### 1.1.2 Retrieval

At retrieval, relational memory is supported by a distributed network including medial temporal, prefrontal, and parietal regions. Within this network, the MTL contributes to reinstatement of relational representations that support recovery of bound episodic features. Prefrontal regions support top-down control processes involved in retrieval search, selection, and monitoring, whereas posterior parietal regions are thought to facilitate the integration of retrieved information into unified episodic representations, potentially functioning as convergence zones for reinstated information (Badre, 2008; Dobbins et al., 2002; Shimamura, 2011).

Despite broad agreement of neural patterns, findings across individual studies remain heterogeneous. Although successful relational retrieval is commonly associated with coordinated engagement of medial temporal, posterior parietal, and prefrontal regions, some studies report particularly robust hippocampal involvement (Nordin et al., 2017; Sullivan Giovanello et al., 2004; Wang & Giovanello, 2016), while others emphasize posterior parietal or prefrontal contributions (Prince et al., 2005; Ranganath et al., 2004; Rugg & Vilberg, 2013), particularly under conditions involving strategic retrieval demands or source monitoring (Dobbins et al., 2002; Wagner et al., 2005).

Variability in task structure, stimulus domain, and the specific contrasts used to isolate relational retrieval success may further contribute to these inconsistencies. As a result, it remains unclear which neural regions are most reliably associated with successful relational retrieval in younger adults across individual studies.

Beyond these unresolved questions in younger adults, neural correlates of relational memory retrieval in aging remain largely heterogeneous and context-dependent. In associative memory paradigms, older adults often recruit a broadly similar cortical retrieval network as younger adults, including medial prefrontal, posterior cingulate, and parietal regions, but show variable hippocampal recruitment and disrupted hippocampal-cortical connectivity (for a review, see Giovanello & Dew, 2015). For example, Wang & Giovanello (2016) reported greater posterior hippocampal activation in older adults when discriminating intact from rearranged word pairs, whereas Tsukiura et al. (2011) observed reduced hippocampal activation in older compared to younger adults during successful retrieval of face–name associations.

These findings suggest that age-related associative memory impairments at retrieval may reflect disruptions in hippocampal-mediated retrieval processes (Hou et al., 2023; Tsukiura et al., 2011).

Similarly, in source memory paradigms, older adults recruit similar core retrieval regions including the hippocampus and parietal cortex, but often exhibit reduced hippocampal engagement, altered hippocampal-cortical connectivity, and decreased recruitment of lateral prefrontal regions that support post-retrieval monitoring compared to younger adults (Cansino et al., 2017; Cansino, Trejo-Morales, et al., 2015; Dulas & Duarte, 2012, 2014). These findings have been interpreted as reflecting age-related disruptions in episodic reinstatement and monitoring processes that support successful source retrieval. At the same time, some studies also report increased engagement of perirhinal or medial prefrontal cortices in older adults, which has also been interpreted as reflecting compensatory or less efficient processing during source memory retrieval (Dulas & Duarte, 2012, 2014). Consequently, it remains unclear which neural regions reliably support successful relational retrieval in older adults and where age-related differences consistently emerge.

Taken together, findings across encoding and retrieval suggest that relational memory in aging is supported by a broadly similar large-scale network, yet the specific pattern of neural engagement within this network varies across studies. Older adults show inconsistent engagement of core regions including the hippocampus, prefrontal cortex, and posterior parietal cortex, with evidence for both increased and decreased recruitment as well as altered hippocampal-cortical interactions. Importantly, this variability appears to depend in part on the cognitive operations emphasized by different relational paradigms, including task demands, relational content, strategic support, and monitoring requirements. Consequently, despite extensive investigation, it remains unclear which regions show reliable convergence across studies and experimental contexts.

### 1.2 Current meta-analysis

As outlined above, extensive research has investigated the neural correlates of relational memory in younger and older adults, yet findings remain difficult to reconcile across studies. Even in younger adults, findings vary across studies, with different studies emphasizing partially distinct medial temporal, prefrontal, and posterior contributions to successful relational memory. Older adults show variable recruitment of core relational memory regions, including the hippocampus and prefrontal cortex, with reports of both age-related increases and decreases in neural engagement during successful relational memory processing. Although these patterns have often been interpreted in terms of compensatory or neural inefficiency (Cabeza et al., 2018; Park & Festini, 2017), such interpretations remain challenging because findings differ across paradigms and experimental contexts. As a result, it remains unclear which neural regions are reliably and consistently engaged during successful relational memory processing within each age group. Coordinate-based meta-analysis provides a critical approach for addressing this issue by identifying regions that show consistent convergence of activation across independent studies while minimizing study-specific variability. By examining convergence separately within each age group and memory phase, such an approach can provide a more stable characterization of the neural networks supporting relational memory and clarify whether core regions, such as the hippocampus, are reliably engaged across younger and older adults.

Although several prior meta-analyses have examined the neural correlates of various memory processes, they have largely focused on younger adults (Kim, 2013, 2020, 2023, 2024a, 2024b, 2025; Kurkela & Dennis, 2016; Spaniol et al., 2009), with one prior study including older adults targeting age-related differences in a broader subsequent memory effect rather than relational memory specifically (Maillet & Rajah, 2014). Despite a substantial behavioral literature and meta-analyses on behavioral data in the domain of associative and relational memory in aging (Fraundorf et al., 2019; Liu et al., 2024; Old & Naveh-Benjamin, 2008; Spencer & Raz, 1995), none to date have examined an equivalent meta-analysis in relational memory within the neuroimaging literature. This gap limits our ability to identify reliable neural patterns associated with relational memory and to understand how these patterns may differ across age groups.

The current meta-analysis aimed to identify convergent neural correlates of relational memory success in younger and older adults using Activation Likelihood Estimation (ALE). We focused specifically on relational memory success during encoding and retrieval, encompassing both associative and source memory paradigms to capture the broader construct of relational binding across different task contexts.

Importantly, we excluded contrasts that compared different types of successful relational binding (e.g., intact pairs vs. rearranged pairs in paired associate paradigms), to isolate neural activity associated with successful relational memory itself, rather than differences between forms of relational processing. Separate meta-analyses were conducted within each age group, as well as direct contrasts between younger and older adults, for each memory phase (encoding and retrieval), enabling characterization of both shared and age-specific neural patterns.

Based on prior work, we hypothesized that core relational memory regions, particularly the hippocampus, would show consistent convergence of activation across both age groups and memory phases, alongside broader engagement of prefrontal regions. However, given the heterogeneity of prior findings, we did not adopt a specific directional hypothesis regarding age-related differences. Instead, we expected the meta-analysis to clarify the extent to which these regions are reliably engaged within each age group and provide a more stable characterization of the neural networks supporting relational memory success.

## 2 Methods

### 2.1 Study selection

To identify studies for inclusion in the current meta-analysis, searches were conducted in three scientific databases (PubMed, PsycINFO, Web of Science; in September 2025). Search terms combined keywords related to relational memory (associative, relational, source, and context), with memory phases (encoding, retrieval) and neuroimaging methods (fMRI, functional MRI). Truncation (e.g., associat, context) was used where appropriate to capture variations in terminology. The full Boolean search string was: (“associat* memory” OR “associat* encoding” OR “associat* retrieval” OR “relational memory” OR “relational encoding” OR “relational retrieval” OR “source memory” OR “source encoding” OR “source retrieval” OR “context* memory” OR “context* encoding” OR “context* retrieval”). The database searches were supplemented by screening the reference lists of relevant review articles on associative memory and neuroimaging (Hwang et al., 2024; Kim, 2023, 2024a; Maillet & Rajah, 2014; Sümer & Kaynak, 2025).

### 2.2 Inclusion criteria

Empirical studies published prior to September 2025 were included if they met the following inclusion criteria:

- Investigated relational memory processes (associative, or source/contextual memory) during encoding and/or retrieval
- Employed fMRI as the primary neuroimaging method
- Included healthy younger and/or older adult participants
- Reported peak activation coordinates from whole-brain analyses, in either Talairach or Montreal Neurological Institute (MNI) space.
- Reported whole-brain activation contrasts within younger and/or older adult groups

### 2.3 Contrast selection

Contrasts of interest included whole-brain, within-age-group contrasts that identified neural correlates of relational memory success. Relational memory success was defined as successful encoding or retrieval of associations between multiple elements of an event (e.g., item-item or item-context relationships), reflecting the binding of those elements into an integrated representation (Opitz, 2010; Ranganath, 2010; Schneegans & Bays, 2017; Yonelinas et al., 2019). For associative memory studies, the current meta-analysis included contrasts of associative hit > associative miss and associative hit > single-item hit. For source memory paradigms, the meta-analysis included contrasts of source hit > source miss, and item + source hit > item-only hit. Collectively, these contrasts enabled us to isolate neural activity associated with successful relational binding, while excluding contrasts that compared different forms of successful relational processing. For example, contrasts such as intact hit > rearranged hit were excluded because correct performance in both conditions can reflect successful relational memory. See the PRISMA flowchart (Figure 1) for full selection procedure.

**Figure 1.**
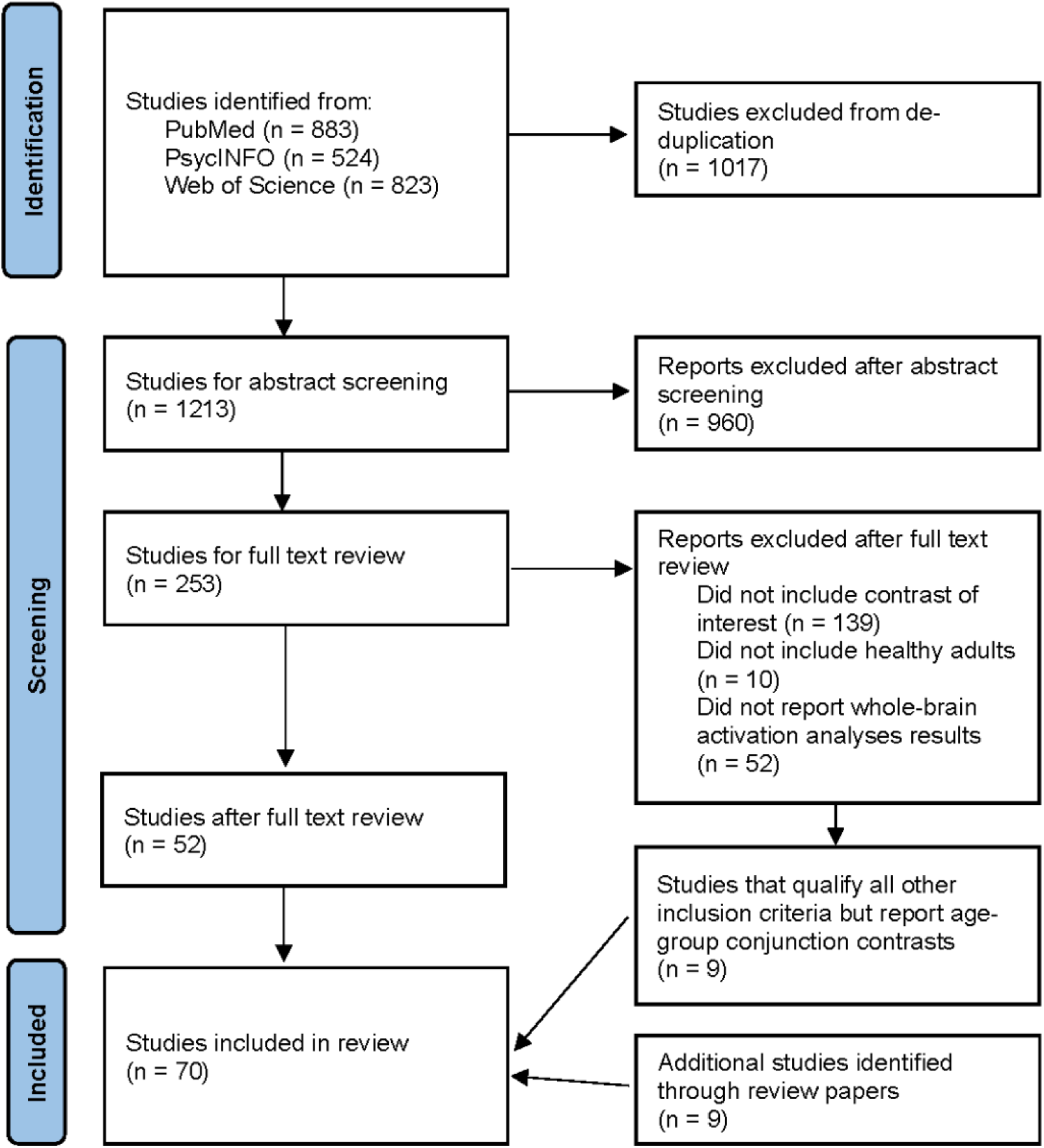
PRISMA flow diagram depicting the study selection process.

The database search yielded 1,213 records after deduplication. Abstract screening excluded 960 records, and full-text review excluded an additional 201 studies, resulting in 52 eligible studies. Nine additional studies were identified through review article screening, and nine studies were included only in the exploratory general relational memory success analysis (see Section 2.4), yielding a total sample of 70 independent studies across all analyses (see PRISMA flow diagram in Figure 1).

### 2.4 Activation likelihood estimation

The current meta-analysis was conducted using the Activation Likelihood Estimation (ALE) method implemented in GingerALE 3.0.2 (www.brainmap.org), following procedures described by Eickhoff and colleagues (Eickhoff et al., 2009, 2012). ALE is a coordinate-based meta-analytic approach that identifies brain regions showing consistent convergence of activation across independent neuroimaging studies.

Reported activation coordinates were modeled as three-dimensional Gaussian probability distributions reflecting spatial uncertainty, which were then combined across experiments to generate voxel-wise ALE scores reflecting the likelihood of convergence of activation foci. Coordinates reported in Talairach space were converted to MNI space using a Lancaster transformation (Lancaster et al., 2007) as implemented in GingerALE 3.0.2. To maintain statistical independence, multiple contrasts derived from the same sample within a study were merged into a single experiment prior to analysis. The resulting ALE map was tested against a null distribution modeling random spatial association of foci across experiments.

Separate meta-analyses were conducted for encoding and retrieval phases within younger and older adults, as well as direct contrasts between age groups for each phase. This design allowed for characterization of both within-group convergence and between-group differences across encoding and retrieval phases. In addition, an exploratory meta-analysis was conducted collapsing across studies, memory phases, and age groups to characterize a general relational memory network. This analysis additionally included conjunction contrasts from studies reporting activation common to younger and older adults (9 studies) but not within-age-group contrasts, as these contrasts reflect activation common to both age groups and are appropriate for analyses that do not distinguish between groups. For within-age-group and exploratory meta-analyses, statistical significance was assessed using a voxel-level cluster-forming threshold of *p* < .001 and a cluster-level familywise error (FWE) threshold of *p* < .05 after 10,000 permutations (as recommended by Eickhoff et al., 2016; Müller et al., 2018). Contrast analyses between age groups were performed using the permutation-based approach implemented in GingerALE (Eickhoff et al., 2011). In this procedure, experiments from both age groups were pooled and randomly reassigned into two groups matching the original sample sizes. ALE maps were recomputed for each permutation, and voxel-wise differences between groups were recorded to generate an empirical null distribution. Observed differences between the original ALE maps were then compared against this null distribution to determine statistical significance.

Significant differences were identified using a voxel-wise threshold of p < .05 with a minimum cluster extent of 500 mm^3^, falling within the range used in prior ALE contrast studies (Bulut, 2023; H. Kim, 2024a, 2025; Papitto et al., 2020). Conjunction analyses reflected voxel-wise overlap between independently thresholded ALE maps from each age group and therefore were not subjected to the permutation-based contrast procedure or cluster extent threshold applied to subtraction analyses. All resulting ALE maps were visualized using the MNI152 template (Fonov et al., 2011) in MRIcroGL (Rorden, 2025a) and Surfice (Rorden, 2025b).

## 3 Results

### 3.1 Encoding

To identify neural regions associated with successful relational memory during encoding, we conducted separate ALE meta-analyses for younger adults, older adults, conjunction analyses across age groups, and direct subtraction analyses comparing younger and older adults.

In younger adults, convergent clusters were observed across frontotemporal regions, including bilateral inferior frontal gyri, the left hippocampus, left parahippocampal/perirhinal cortices, and bilateral fusiform gyri. In contrast, older adults showed more limited convergence, with clusters observed in the left hippocampus and right inferior temporal gyrus.

Conjunction analyses revealed shared convergence between age groups in the left hippocampus and right inferior temporal gyrus. Direct contrasts between age groups further revealed greater convergence in younger relative to older adults in the left inferior frontal gyrus, with no regions showing greater convergence in older relative to younger adults. Full results are reported in Table 1 and Figure 2.

**Figure 2.**
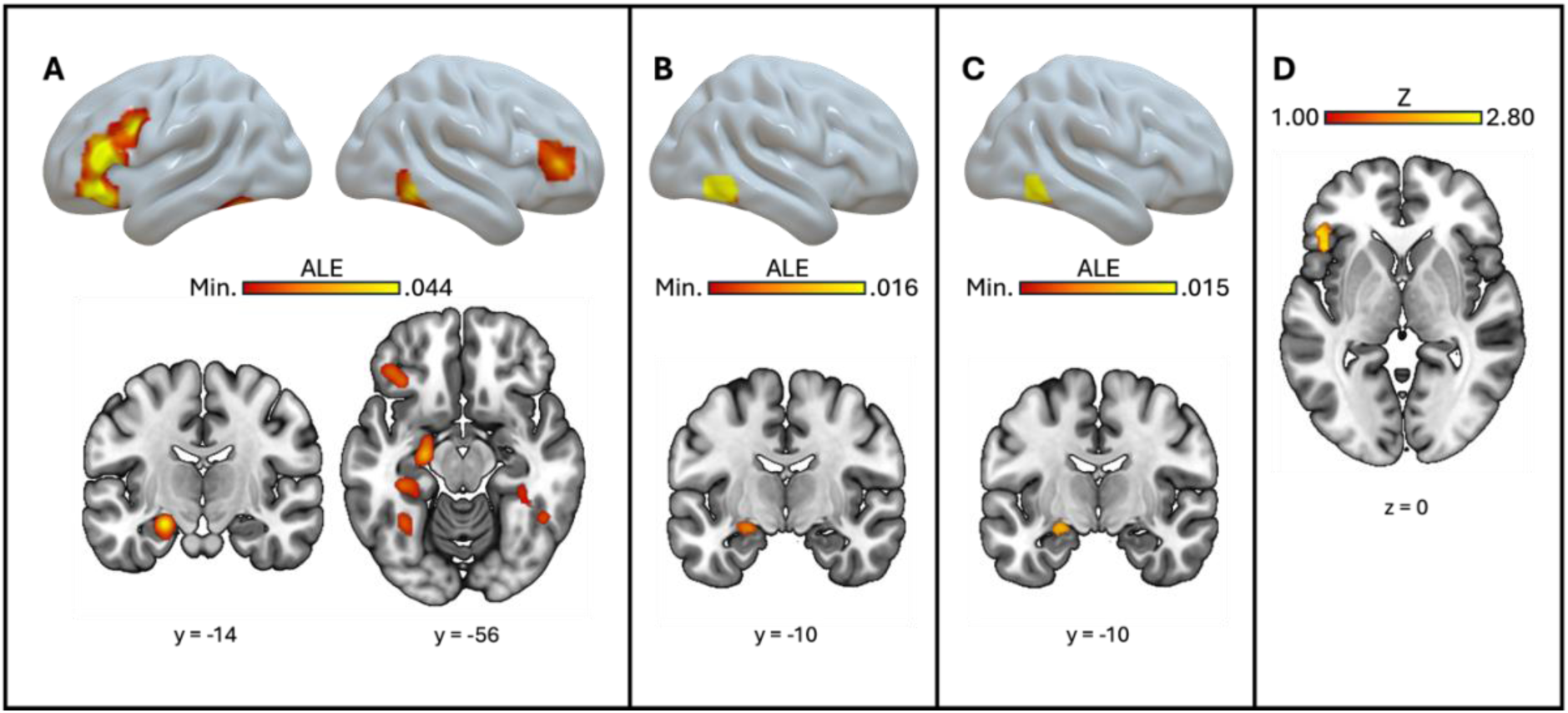
Regions associated with relational memory success during encoding. (A) Within younger adults, (B) within older adults, (C) across both age groups, and (D) younger > older.

**Table 1.** Results of ALE meta-analyses for relational memory success during encoding within, across, and between age groups.

| Encoding | Region | *H* | Peak MNI |  |  |  | *Z* | Volume (mm<sup>3</sup>) |
| --- | --- | --- | --- | --- | --- | --- | --- | --- |
|  |  |  | *x* | *y* | *z* | ALE |  |  |
| Younger | Inferior Frontal Gyrus | L | -48 | 28 | 12 | .037 | 6.31 | 6672 |
| (456 foci, 42 exp) | Hippocampus | L | -22 | -14 | -16 | .044 | 7.00 | 2824 |
|  | Inferior Frontal gyrus | L | -42 | 8 | 30 | .024 | 4.62 | 1936 |
|  | Parahippocampal/<br>Perirhinal Cortex | L | -34 | -34 | -12 | .028 | 5.13 | 1800 |
|  | Fusiform Gyrus | L | -34 | -56 | -16 | .026 | 4.92 | 1376 |
|  | Inferior Temporal Gyrus | R | 48 | -52 | -10 | .029 | 5.30 | 1088 |
|  | Fusiform Gyrus | R | 34 | -36 | -12 | .021 | 4.21 | 1032 |
|  | Dorsolateral Prefrontal<br>Cortex | R | 44 | 36 | 6 | .021 | 4.25 | 1024 |
| Older | Inferior Temporal Gyrus | R | 52 | -58 | -10 | .016 | 4.81 | 720 |
| (85 foci, 11 exp) | Hippocampus | L | -22 | -10 | -14 | .011 | 3.77 | 616 |
| Conjunction | Hippocampus | L | -22 | -10 | -14 | .011 | - | 504 |
|  | Inferior Temporal Gyrus | R | 52 | -56 | -10 | .015 | - | 224 |
| Younger > Older | Inferior Frontal Gyrus | L | -44 | 20 | 0 | - | 2.79 | 2808 |
| Older > Younger | - |  |  |  |  |  |  |  |

**Note.** H = hemisphere; x, y, and z denote peak MNI coordinates; ALE = activation likelihood estimation value at the peak voxel; Z = Z score associated with the reported ALE or contrast statistic.

### 3.2 Retrieval

To identify neural regions associated with successful relational memory during retrieval, we conducted separate ALE meta-analyses for younger adults, older adults, conjunction across both age groups, and direct subtraction analyses contrasting younger and older adults.

In younger adults, convergent clusters were observed in posterior midline and subcortical regions, including the posterior cingulate/precuneus, thalamus, and bilateral putamen. In contrast, older adults showed one significant cluster in the left angular gyrus.

Conjunction analyses revealed no shared convergence between age groups. Direct contrasts between age groups revealed greater convergence in younger relative to older adults in the posterior cingulate cortex, with no regions showing greater convergence in older relative to younger adults. Full results are reported in Table 2 and Figure 3.

**Figure 3.**
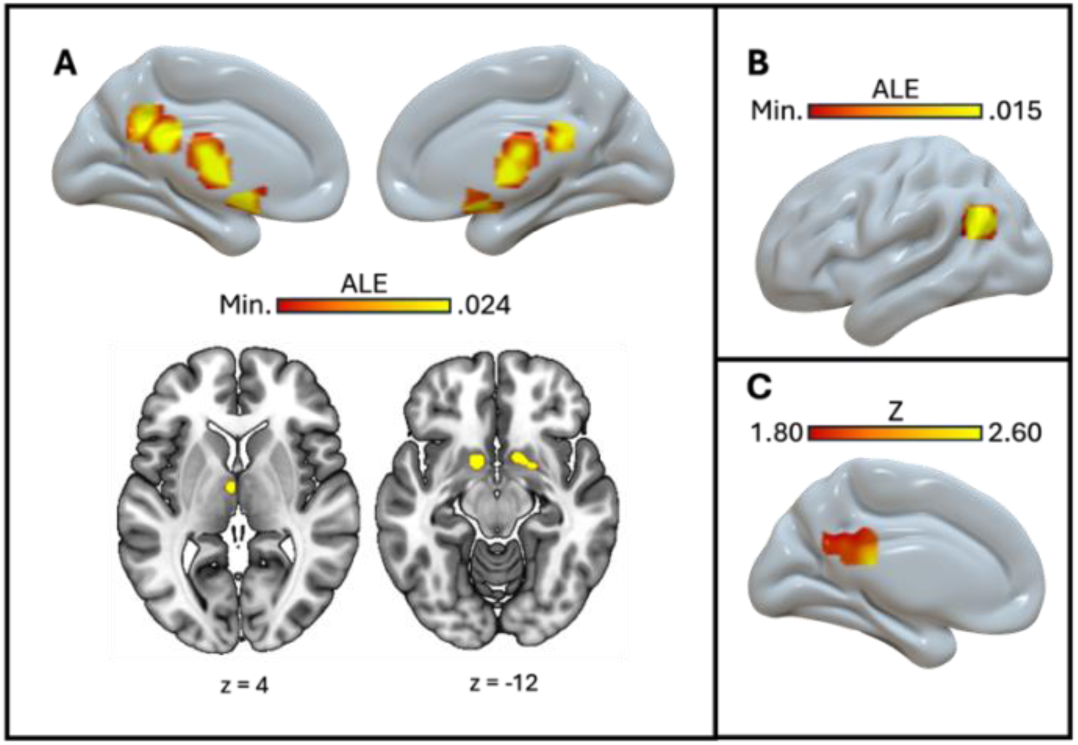
Regions associated with relational memory success during retrieval. Within younger adults (A), within older adults (B), and younger > older (C).

**Table 2.**
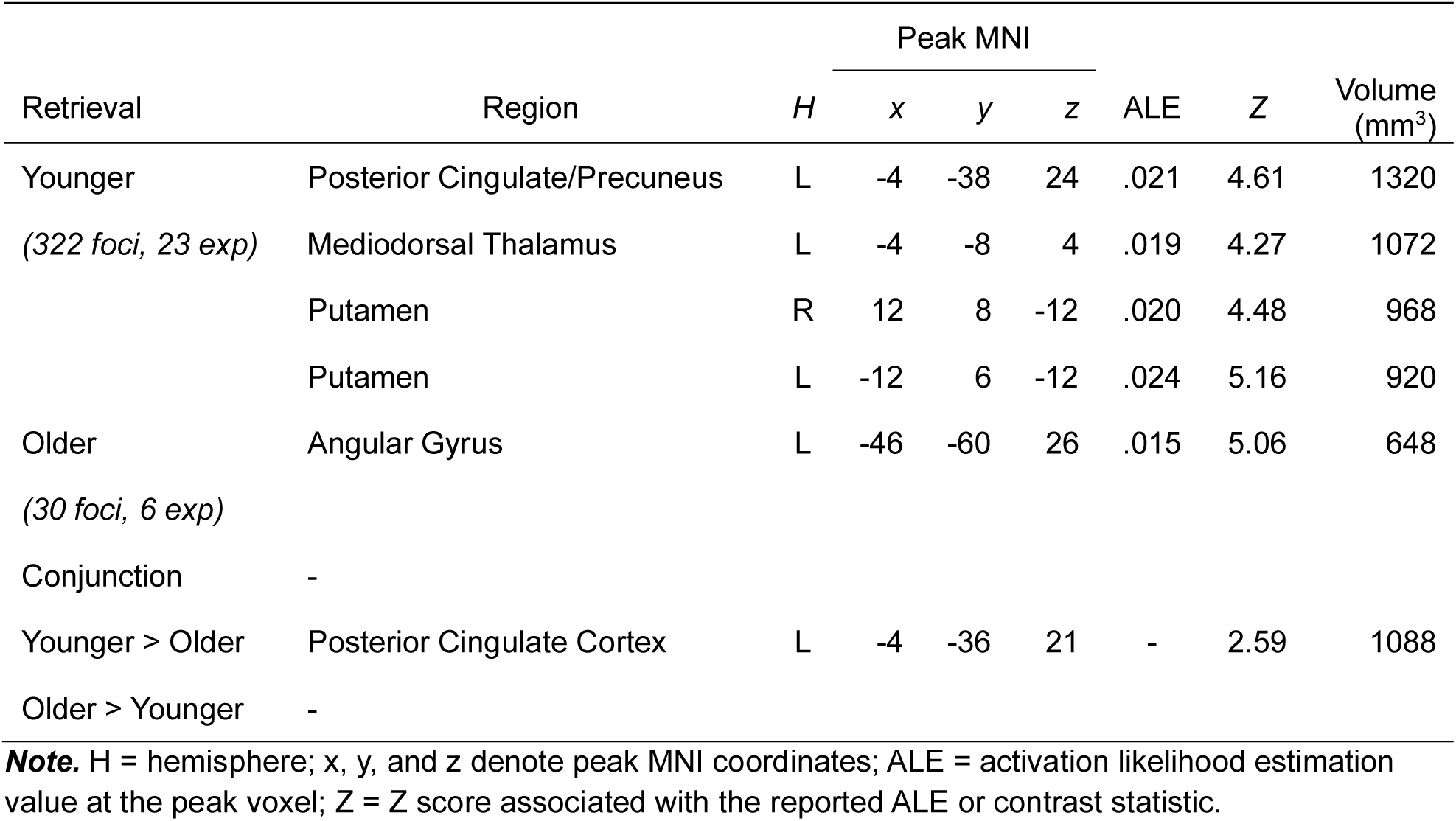
Results of ALE meta-analyses for relational memory success during retrieval within, across, and between age groups.

### 3.3 General relational memory network

To characterize a general relational memory success network, we conducted an exploratory ALE meta-analysis collapsing across all studies and memory phases (encoding and retrieval), with the addition of studies that reported conjunction contrasts but no within-age contrasts. Significant convergence was observed across medial temporal, prefrontal, posterior parietal, and occipitotemporal regions. Full results are reported in Table 3 and Figure 4.

**Figure 4.**
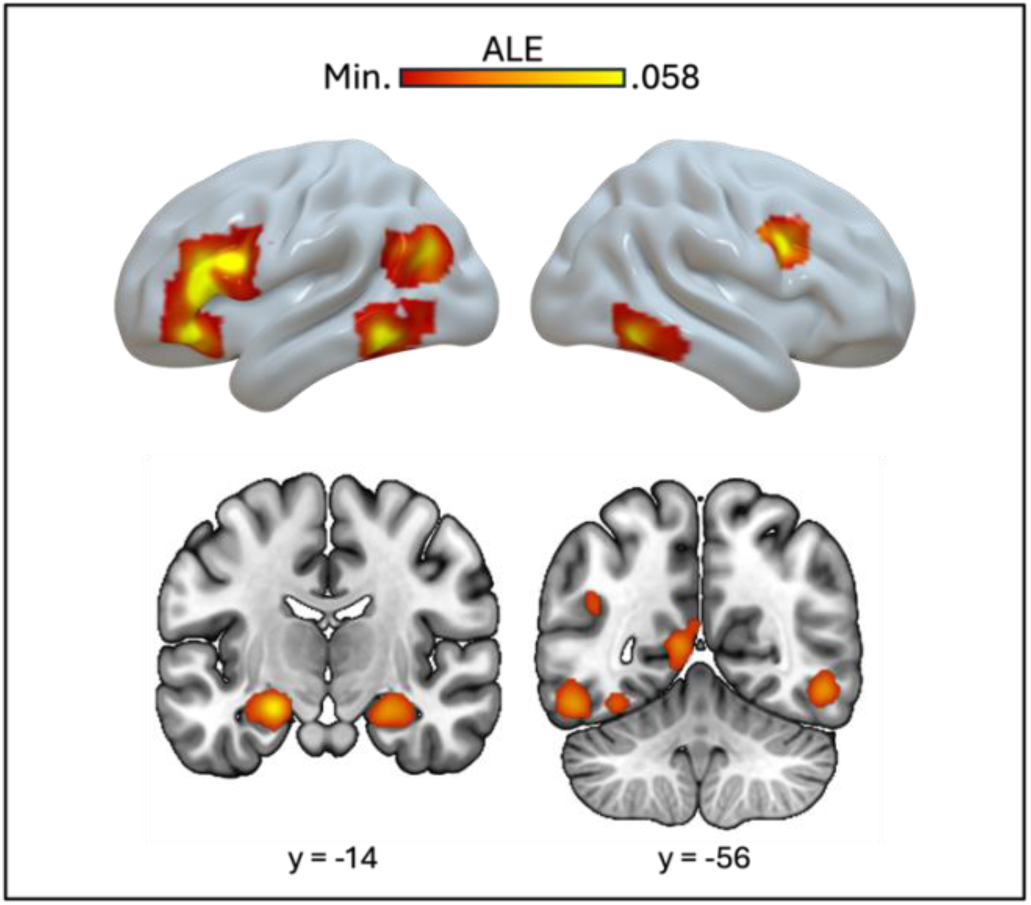
Regions associated with relational memory success in general. Regions showing significant convergence across all included relational memory studies collapsed across encoding and retrieval phases and across age groups.

**Table 3.** Results of ALE meta-analyses for relational memory across all studies.

| Region | *H* | Peak MNI |  |  | ALE | *Z* | Volume<br>(mm<sup>3</sup>) | Contributing<br>Experiments |
| --- | --- | --- | --- | --- | --- | --- | --- | --- |
|  |  | *x* | *y* | *z* |  |  |  |  |
| Inferior Frontal Gyrus | L | -50 | 28 | 10 | .052 | 6.86 | 11768 | 45 |
|  |  |  |  |  | (28 *yENC*; 9 *yRET*; 3 *oENC*; 3 *cENC*; 3 *cRET*) |  |  |  |
| Hippocampus | L | -22 | -14 | -16 | .058 | 7.38 | 8336 | 35 |
|  |  |  |  |  | (18 *yENC*; 8 *yRET*; 4 *oENC*; 3 *cENC*; 2 *cRET*) |  |  |  |
| Hippocampus | R | 26 | -10 | -20 | .043 | 5.90 | 4632 | 25 |
|  |  |  |  |  | (10 *yENC*; 7 *yRET*; 4 *oENC*; 1 *oRET*; 2 *cENC*; 1 *cRET*) |  |  |  |
| Fusiform Gyrus | L | -52 | -54 | -16 | .042 | 5.78 | 4384 | 24 |
|  |  |  |  |  | (10 *yENC*; 1 *yRET*; 2 *oENC*; 3 *cENC*; 2 *cRET*) |  |  |  |
| MTG/Angular Gyrus | L | -44 | -60 | 26 | .035 | 5.04 | 2840 | 15 |
|  |  |  |  |  | (6 *yENC*; 4 *yRET*; 1 *oENC*; 1 *oRET*; 2 *cENC*; 1 *cRET*) |  |  |  |
| Posterior Cingulate Cortex | L | -8 | -56 | 8 | .038 | 5.38 | 2256 | 10 |
|  |  |  |  |  | (4 *yENC*; 2 *yRET*; 1 *oENC*; 2 *cENC*; 1 *cRET*) |  |  |  |
| Fusiform Gyrus | R | 50 | -54 | -10 | .043 | 5.92 | 2104 | 14 |
|  |  |  |  |  | (8 *yENC*; 2 *yRET*; 2 *oENC*; 1 *cENC*; 1 *cRET*) |  |  |  |
| Precentral Gyrus | R | 46 | 8 | 30 | .044 | 6.01 | 1232 | 7 |
|  |  |  |  |  | (3 *yENC*; 1 *oENC*; 1 *oRET*; 1 *cENC*; 1 *cRET*) |  |  |  |

**Note.** *H* = hemisphere; *x*, *y*, and *z* denote peak MNI coordinates; ALE = activation likelihood estimation value at the peak voxel; *Z* = *Z* score associated with the reported ALE statistic. This exploratory analysis additionally included conjunction contrasts reporting activation common to younger and older adults. *yENC/yRET* denote younger adult encoding/retrieval experiments, *oENC/oRET* denote older adult encoding/retrieval experiments, and *cENC/cRET* denote experiments reporting conjunction contrasts for encoding/retrieval, respectively.

## 4 Discussion

The present meta-analysis provides the first quantitative characterization of the neural correlates of relational memory success across younger and older adults. Across encoding and retrieval, several consistent patterns emerged. During encoding, both age groups showed convergence in the hippocampus and inferior temporal gyrus. Younger adults additionally showed convergence across prefrontal and other medial temporal regions, whereas older adults showed no additional convergence. During retrieval, younger adults showed convergence in posterior midline and subcortical regions, whereas older adults showed convergence in the angular gyrus, with no shared convergence between age groups. Across all analyses, the hippocampus showed the most consistent bilateral convergence. Findings are discussed first by memory phase, followed by consideration of the broader relational memory network and implications for age-related differences.

### 4.1 Relational memory during encoding

Relational memory encoding requires the integration of multiple event elements into a unified representation that can support subsequent retrieval. Consistent with this demand, the present meta-analysis revealed a distributed set of regions including medial temporal, prefrontal, and ventral temporal regions supporting relational encoding. Among these regions, the left hippocampus emerged as the most robust and consistent region of convergence across both younger and older adults.

In addition to being observed within each age group, the left hippocampus was the only MTL region shared across age groups in the conjunction analysis. This finding is especially noteworthy given the heterogeneity of paradigms and materials contributing to it, including associative memory tasks involving word-word, face-name, face-scene, and object-location pairings, as well as source memory paradigms requiring the encoding of spatial, temporal, and perceptual contextual features. Despite differences in task structure and stimulus content, these paradigms share a common computational demand: multiple event elements must be integrated into a single relational representation. Thus, the present meta-analytic convergence suggests that hippocampal recruitment represents a reliable and domain-general signature of relational encoding across diverse forms of associative and contextual information (Olsen et al., 2012; Park et al., 2012).

This finding is consistent with prior work proposing that the hippocampus serves as a convergence zone that binds item and context information provided by surrounding MTL cortices into an integrated episodic representation while preserving relations among event features (Davachi, 2006; Konkel & Cohen, 2009; Libby et al., 2019; Sullivan Giovanello et al., 2004; Yonelinas et al., 2019). Importantly, this role appears to be defined less by the specific content being encoded than by the relational computation required. Converging evidence from single unit recordings and patient populations supports this view. Human single-neuron studies indicate that hippocampal neurons bind items to their spatiotemporal context and support the reinstatement of those bound representations (Kunz et al., 2024; Miller et al., 2013), and that hippocampal activity correlates with associative retrieval success independent of item identity (Rutishauser et al., 2021). Patient and fMRI studies suggest that the hippocampus is particularly critical when associations must be formed across stimulus domains, supporting integration of distinct representational elements into a unified memory trace (Borders et al., 2017; Mayes et al., 2007; Mayes et al., 2004; Piekema et al., 2009; Vargha-Khadem et al., 1997). Notably, the presence of this convergence in both younger and older adults suggests that hippocampal engagement represents one of the most stable neural correlates of relational encoding across the adult lifespan (Giovanello & Dew, 2015).

In addition to the hippocampus, both younger and older adults showed convergence in the right inferior temporal gyrus (ITG). The ITG is a core region of the ventral visual stream and is implicated in high-level object processing and representation of visual features (Conway, 2018; Martin & Barense, 2023; Nobre et al., 1994). Its consistent recruitment across age groups suggests that successful relational encoding relies, in part, on the availability of detailed item-level representations within the ventral visual stream itself. This finding is consistent with representational-hierarchical accounts, which propose that ventral temporal regions, including the ITG, form a continuous processing hierarchy organized by representational complexity (Martin & Barense, 2023). Within this framework, ITG representations support relational encoding by providing the high-level feature conjunctions necessary for linking items with their contextual associations. Consistent with this view, prior work shows that activity in ventral temporal cortex reflects the content of information maintained or reactivated in memory (Ranganath et al., 2004), and meta-analytic evidence suggests that the ITG contributes to identifying the item or event to which contextual information is bound in episodic memory (Torres-Morales & Cansino, 2024). The presence of this convergence in both younger and older adults suggests that access to detailed item-level representations is a shared component of relational encoding across the adult lifespan.

Younger adults showed additional convergence in the left parahippocampal/perirhinal cortex. Within the MTL, the perirhinal cortex (PRC) is thought to support item-level representation and familiarity-based processing, whereas the parahippocampal cortex (PHC) supports encoding of contextual and spatial information (Aminoff et al., 2013; Diana et al., 2007; Suzuki & Naya, 2014). The experiments contributing to this cluster place varying demands on these representational systems. For example, associative tasks involving word–word or face–name pairs rely more heavily on item-level and semantic representations, whereas object–location and source memory paradigms place greater demands on contextual and spatial encoding. Despite this variation, convergence within a cluster encompassing both the PRC and PHC suggests that successful relational encoding engages both item-level and contextual representations, even when one type of information is more task-relevant than the other. In the present meta-analysis, this pattern was observed in younger adults, suggesting that successful relational encoding may depend on the concurrent availability of both item and contextual information, highlighting the interdependence of these representations in episodic memory formation (Park et al., 2012).

Beyond the MTL, younger adults also showed robust convergence across prefrontal regions, including two clusters in the left inferior frontal gyrus (IFG) and a right dorsolateral prefrontal cortex (DLPFC). The two left IFG clusters map onto functionally distinct subregions identified in prior meta-analytic work (Liakakis et al., 2011). The larger, more anterior cluster (MNI: −48, 28, 12) corresponds to a region associated with semantic and phonological processing, whereas the more dorsal cluster (MNI: −42, 8, 30) falls within a region associated with verbal working memory (Liakakis et al., 2011).

Alongside prior work highlighting the role of the left IFG in semantic and verbal processing (Chou et al., 2009; Huang et al., 2012; Roskies et al., 2001), these findings suggest that the IFG supports at least two complementary processes during relational encoding: semantic integration of to-be-bound elements, and the maintenance of those elements in working memory during relational binding. Supporting this interpretation, meta-analytic evidence implicates the IFG in paired associate recollection (Kim, 2023), source memory (Kim, 2020, 2025), and associative word encoding (Kim, 2024a). The right DLPFC similarly supports the organization and monitoring of multiple elements in working memory during encoding. Prior work suggests that this region contributes to long-term memory formation by strengthening associations among actively maintained and monitored representations (Balconi, 2013; Blumenfeld et al., 2011; Blumenfeld & Ranganath, 2006; Melrose et al., 2020; Sandrini et al., 2003).

Together, these findings suggest that successful relational encoding is supported by core mechanisms including hippocampal binding and ventral temporal representations in the ITG. In younger adults, this core system is further complemented by additional medial temporal contributions from the perirhinal and parahippocampal cortices as well as prefrontal processes that support the integration and maintenance of to-be-bound information. Although these additional regions did not emerge as significant in older adults, this pattern should be interpreted cautiously given the smaller number of older adult encoding studies included in the meta-analysis. Broader implications of these age-related differences are considered further in Section 4.4.

### 4.2 Relational memory success during retrieval

Relational memory retrieval was supported by a distinct pattern of neural convergence in each age group, with younger adults showing convergence in posterior midline and subcortical regions and older adults showing convergence in the angular gyrus. In younger adults, the posterior cingulate cortex (PCC)/precuneus emerged as the most robust region of convergence. These regions have previously been implicated in episodic memory retrieval and are thought to support the recovery of contextual and relational information associated with past experiences (Dadario & Sughrue, 2023; B. L. Foster & Koslov, 2025). For example, PCC activity has been shown to increase with the subjective strength and contextual detail of retrieved memories, with its ventral subregion specifically implicated in effortful recollection and the retrieval of spatial and contextual features (Foster et al., 2023). Similarly, the precuneus has been linked to source memory retrieval and the reinstatement of contextual associations, such as when and where items were previously encountered (Lundstrom et al., 2003, 2005).

Together, these findings suggest that relational memory retrieval in younger adults is supported by posterior midline regions that contribute to the recovery and representation of contextual information. Additional convergence in younger adults was observed in the mediodorsal thalamus and bilateral putamen. The mediodorsal thalamus maintains dense structural connections with the prefrontal cortex and is believed to support memory retrieval by maintaining and coordinating activity within frontal networks (Mitchell & Chakraborty, 2013; Pergola et al., 2018). The bilateral putamen convergence may reflect processes related to successful retrieval outcomes, consistent with the role of the striatum as a core component of the brain’s reward system (Cromwell et al., 2005; Schultz, 2000), as well as prior work showing that striatal activity during episodic retrieval is associated with the successful recovery of information and the subjective satisfaction of recollection (Dong et al., 2016; Shigemune et al., 2017; Weinstein, 2023). Supporting this account, prior meta-analytic evidence indicates stronger convergence in the putamen for contrasts isolating successful retrieval (e.g., hit > miss), compared to contrasts such as memory > perception, which include both successful and unsuccessful trials (Kim et al., 2023). Similarly, the present meta-analysis included only contrasts indicative of successful retrieval, further suggesting that putamen convergence reflects processes specifically associated with successful recollection.

In older adults, convergence was limited to a single cluster in the left angular gyrus. The angular gyrus has been consistently implicated in episodic memory retrieval and is considered a core node of the default network (Bonnici et al., 2016; Humphreys et al., 2021). Prior studies suggest a specific role for this region in detailed episodic recollection, with activity associated with more vivid and specific mnemonic detail (Bellana et al., 2016; Bonnici et al., 2016; Thakral et al., 2017). Of note, the two experiments contributing to this cluster used similar object-location paradigms (Cansino et al., 2015; Cansino, et al., 2017).

In contrast to the robust MTL convergence observed during encoding, no cluster within the MTL emerged as significant during relational memory retrieval (but see Section 4.3 for evidence of MTL convergence in the collapsed analysis). This absence should be interpreted cautiously. One important consideration is statistical power, as fewer studies were included in the retrieval analyses overall, with particularly limited representation of older adult retrieval studies. This issue is discussed in greater detail in the Limitations section.

In addition to power, characteristics of the contrasts used in the present meta-analysis may also contribute to the absence of MTL convergence. Although both encoding and retrieval analyses relied on success-based contrasts, the nature of these contrasts differs across memory phases. During encoding, subsequent memory contrasts (e.g., remembered > forgotten) typically capture hippocampal activity that is more closely associated with successful memory formation, resulting in a strong signal that survives subtraction (Cansino et al., 2015; Halpern et al., 2023; Maillet & Rajah, 2014; Park et al., 2013). In contrast, during retrieval, contrasts such as hit > miss may attenuate hippocampal effects if the hippocampus is engaged during both successful and unsuccessful retrieval attempts, including processes such as cue-driven reactivation or partial retrieval (Doshier & Ryals, 2020; Frankland et al., 2019). In such cases, subtraction-based contrasts may reduce the magnitude of hippocampal activation, resulting in less consistent convergence across studies. Consistent with this interpretation, prior meta-analytic work has reported less consistent hippocampal convergence during retrieval relative to encoding, particularly for contrasts isolating retrieval success, whereas contrasts emphasizing relational comparison or mismatch detection (e.g., intact vs. rearranged pairs) show more robust hippocampal convergence (Kim, 2023; Spaniol et al., 2009).

Taken together, the absence of MTL convergence in the present retrieval analyses likely reflects a combination of reduced statistical power and properties of the contrast structure, rather than a lack of hippocampal involvement in relational memory retrieval.

### 4.3 General relational memory success

The exploratory meta-analysis collapsing across all studies, phases, and age groups revealed a broad network of regions consistently engaged during relational memory success. This network includes medial temporal, prefrontal, posterior parietal, and occipitotemporal regions, and closely mirrors the broader episodic memory network identified in prior research (Kim, 2013, 2023; Rugg & Vilberg, 2013; Spaniol et al., 2009). Importantly, these findings suggest that relational memory success is supported by a core, domain-general network that is consistently recruited across task contexts, memory phases, and age groups.

Within this general network, the bilateral hippocampi emerged as robust regions of convergence. Contributing experiments for both left and right hippocampal clusters were distributed across both encoding and retrieval phases, and across both age groups (see Table 3), emphasizing the critical role of the hippocampus in relational memory success, regardless of phase or age. Notably, hippocampal convergence in the general meta-analysis suggests that the lack of hippocampal convergence in the retrieval-specific analyses does not reflect a genuine absence of hippocampal involvement during relational memory retrieval. Rather, as noted above, this pattern likely reflects reduced power in retrieval analyses due to the smaller number of retrieval studies relative to encoding studies. This issue may have been further exacerbated by our inclusion criteria. Specifically, a substantial number of studies excluded from the meta-analysis due to the absence of whole-brain analyses (24 studies) reported significant hippocampal activation associated with relational memory success, using ROI-based approaches. Taken together, the robust hippocampal convergence observed across the relational memory network supports established accounts of the hippocampus as a critical region for relational binding regardless of phase and age (Giovanello & Dew, 2015; Konkel & Cohen, 2009).

The left IFG also showed a large convergent cluster in the general relational memory analysis. As discussed above, the left IFG likely supports semantic integration of to-be-bound elements and the maintenance of those elements in working memory during relational binding. This convergence is consistent with prior work implicating the

IFG in controlled semantic processing and the organization of information during memory formation. Together, these findings suggest that the IFG contributes to relational memory success by supporting integration and maintenance of multiple representations necessary for successful binding.

Lastly, the bilateral fusiform gyri showed convergence in the general relational memory network. The fusiform gyrus is associated with high-level visual and object processing (Weiner & Zilles, 2016; Zhang et al., 2016), and its bilateral convergence here likely reflects the perceptual processing of stimuli across the range of relational memory paradigms included in the meta-analysis. Specifically, the fusiform gyrus has been implicated in the processing of faces (Kanwisher & Yovel, 2006; McCarthy et al., 1997; Rossion et al., 2024), visual words (Dehaene et al., 2002; Devlin et al., 2006; McCandliss et al., 2003), and objects (Starrfelt & Gerlach, 2007; Weiner et al., 2018) — stimulus categories that are well represented across the experiments contributing to these clusters.

### 4.4 Age-related differences in relational memory

Interpretation of age-related differences in the present meta-analysis should be made with caution. Although the statistical comparisons between age groups were conducted using stringent permutation-based approaches, the number of older adult experiments was substantially smaller than that of younger adult experiments across both encoding and retrieval analyses. This imbalance may limit the representativeness and stability of convergence observed in older adults, particularly for null findings. As such, age-related differences should be interpreted with the consideration that asymmetries in the underlying literature may influence the pattern of observed convergence.

Direct comparisons between age groups revealed greater convergence in younger relative to older adults in the left IFG during encoding and in the PCC during retrieval. The IFG finding is consistent with age-related differences in the engagement of controlled semantic and strategic processes during relational encoding, as this region has been associated with semantic elaboration, controlled retrieval, and the organization of information during encoding (Craik, 2023; Glisky, 2002). Similarly, greater convergence in younger adults in the PCC during retrieval suggests more consistent engagement of posterior-medial processes supporting the representation and evaluation of retrieved episodic content. The PCC is a core component of posterior-medial memory systems and has been implicated in integrating retrieved information into coherent representations that guide behavioral responses (Foster et al., 2023; Rugg & Vilberg, 2013).

In addition to these direct effects, qualitative differences in within-group convergence provide further context for interpreting age-related patterns. During encoding, convergence in parahippocampal and perirhinal cortices was observed in younger but not older adults. This asymmetry may reflect a pattern consistent with differences in the availability or reliability of item and contextual representations that support relational binding. Successful relational memory depends not only on hippocampal binding processes but also on the availability of distinct item and contextual information supported by surrounding medial temporal regions. Prior work suggests that aging is associated with reduced representational specificity in ventral temporal and medial temporal cortices (Koen & Rugg, 2019; Park & Festini, 2017), which may impact the fidelity of information available for binding. However, because the direct younger > older contrast did not reveal a significant difference in these regions, this pattern should not be interpreted as definitive evidence of reduced medial temporal recruitment in older adults, but rather as a qualitative difference that may reflect variability in convergence across studies.

Taken together, these findings indicate that age-related differences in relational memory are most consistently observed in regions supporting controlled encoding processes and the evaluation of retrieved information, as indexed by greater convergence in younger adults in the IFG and PCC. At the same time, qualitative differences in within-group convergence, particularly within medial temporal regions, provide additional context for interpreting these patterns but do not constitute direct evidence of age-related differences. Given the imbalance in the underlying literature, these findings should be interpreted as provisional, highlighting the need for future work with more balanced samples to more precisely characterize age-related differences in the neural mechanisms supporting relational memory.

## 5 Limitations and Conclusions

Several limitations of the current meta-analysis should be considered when interpreting the findings. The most important limitation concerns the imbalance in the number of experiments available for younger versus older adults. Older adult experiments were substantially underrepresented across both encoding (OA:11; YA:42) and retrieval (OA:6; YA:23) analyses. This imbalance may limit the representativeness of convergence observed in older adults, particularly for null findings. As such, age-related differences should be interpreted with the understanding that asymmetries in the included studies may influence the observed patterns.

A related consideration concerns the absence of MTL convergence during retrieval. As discussed above, this pattern likely reflects a combination of reduced statistical power and characteristics of the contrasts included in the meta-analysis, rather than a true absence of hippocampal involvement in successful relational memory retrieval. Consistent with this interpretation, hippocampal convergence was observed in the collapsed analysis across phases, and many studies excluded from the present meta-analysis due to the absence of whole-brain analyses reported significant MTL involvement using ROI approaches. Together, these observations suggest that the lack of MTL convergence during retrieval should be interpreted as a methodological constraint rather than a functional absence.

More broadly, as with all coordinate-based meta-analyses, the present findings are influenced by variability in MRI acquisition parameters, preprocessing pipelines, and statistical thresholds across included studies, introducing heterogeneity that cannot be fully controlled. Additionally, ALE identifies regions of spatial convergence, but does not capture functional connectivity or directly assess brain-behavior relationships. As such, interpretations of identified clusters rely on convergence with the broader neuroimaging literature rather than direct inference from the meta-analytic data alone.

Despite these limitations, the present study provides the first quantitative synthesis of the neural correlates of relational memory success in younger and older adults. Across encoding and retrieval phases, the hippocampus emerged as one of the most consistent and robust regions of convergence, underscoring its central role in relational binding across the adult lifespan. Beyond this core contribution, relational memory success was supported by a broader network spanning medial temporal, prefrontal, posterior parietal, and occipitotemporal regions, reflecting the integration of binding, control, and representational processes.

At the same time, the findings highlight variability in the engagement of this network across age groups, particularly in regions supporting controlled encoding and the evaluation of retrieved information. Importantly, these patterns should be interpreted in the context of the current literature, which remains heavily weighted toward younger adult samples. Together, these results provide a quantitative foundation for understanding the neural correlates of relational memory success and identify both stable and variable components of the relational memory success network across the adult lifespan. They further emphasize the need for greater representation of older adults in neuroimaging research to more precisely characterize how relational memory processes change with age.

## Data and Code Availability

The ALE maps resulting from all meta-analyses in this study are openly available on NeuroVault (https://identifiers.org/neurovault.collection:23735).

## Author Contributions

MS: conceptualization, methodology, software, validation, formal analysis, investigation, data curation, writing - original draft, writing - review & editing, visualization, project administration. ND: conceptualization, validation, resources, writing-review & editing, supervision.

## Supplementary Material

**Supplementary Table 1.**
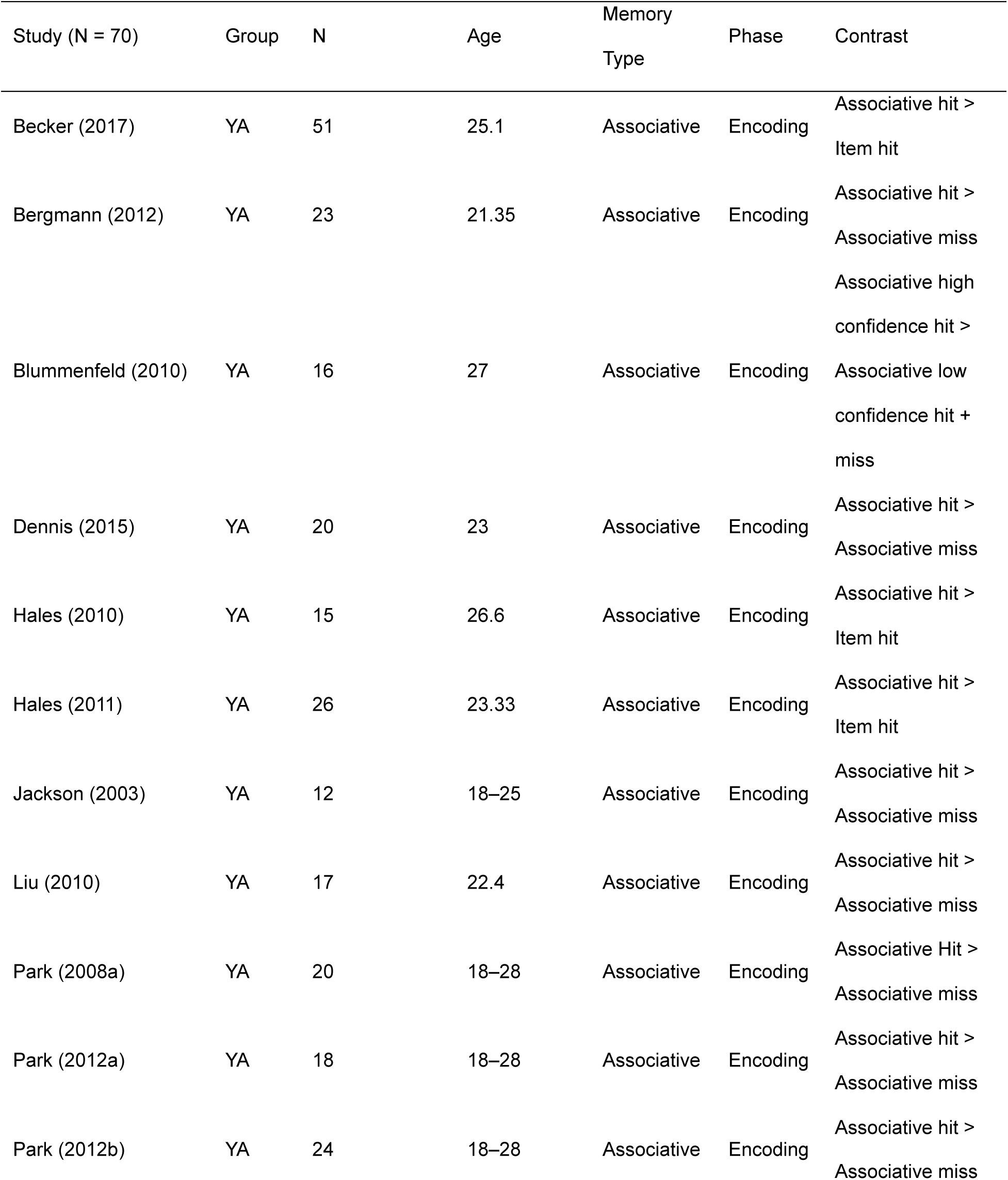
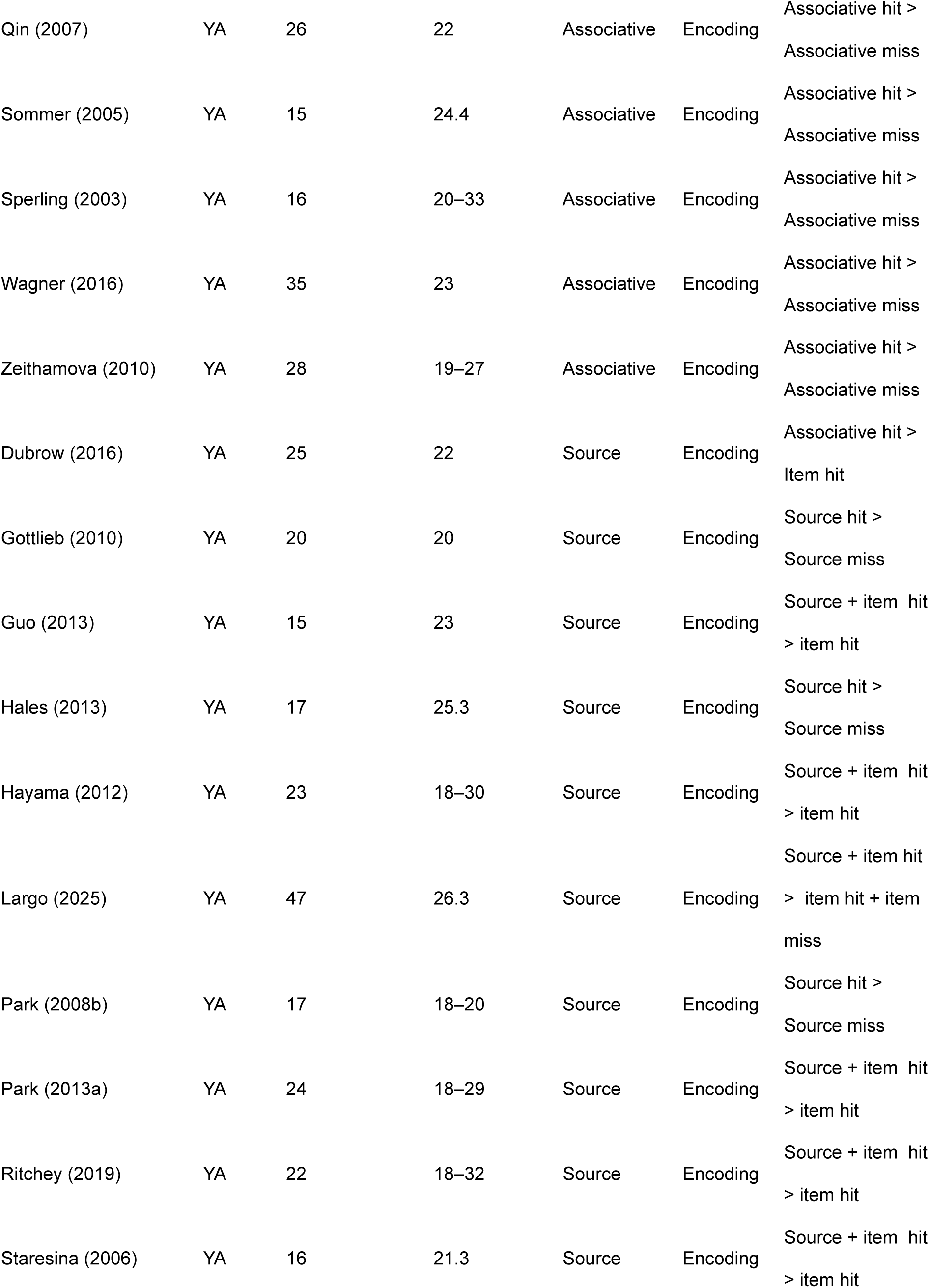
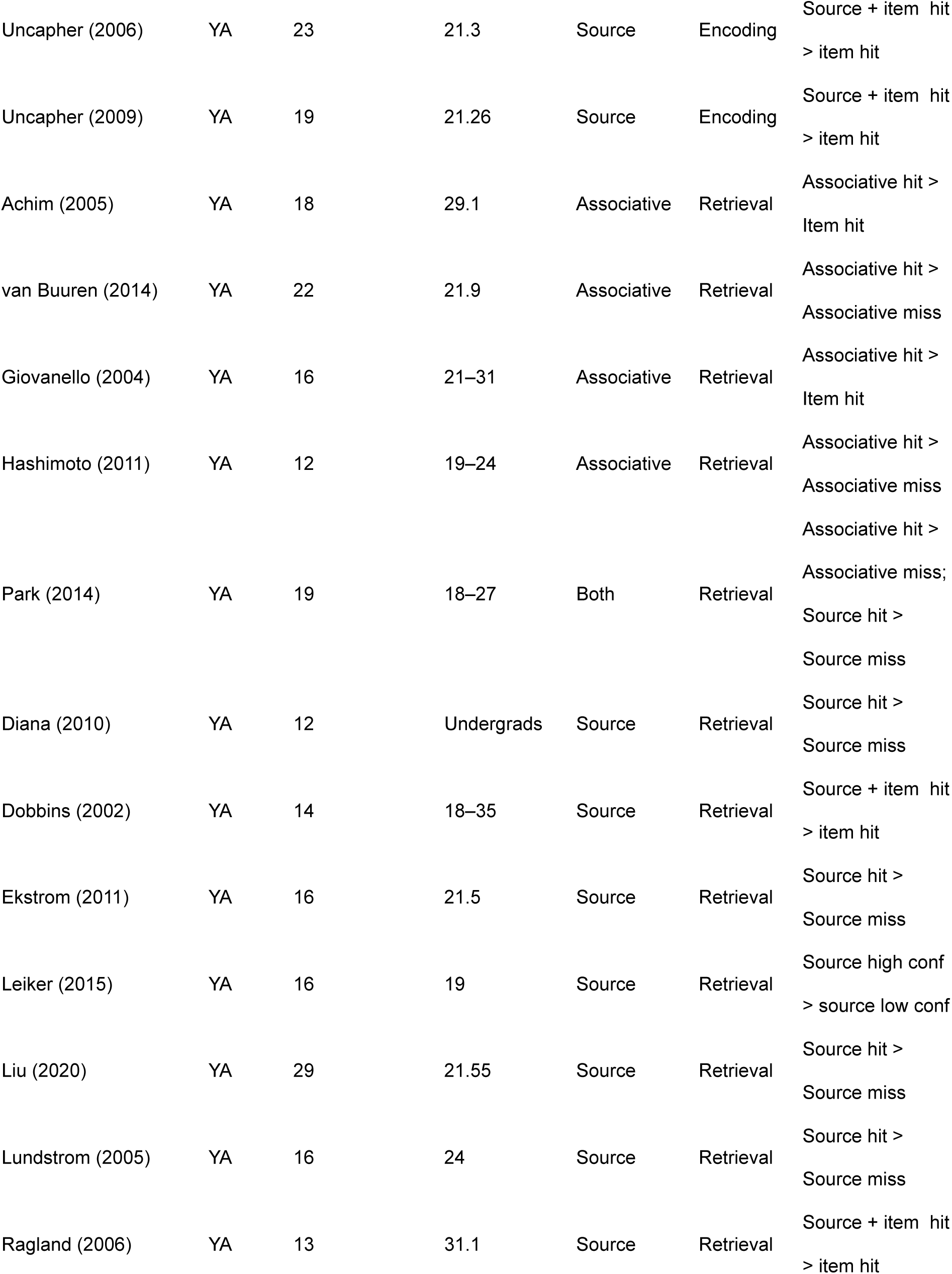
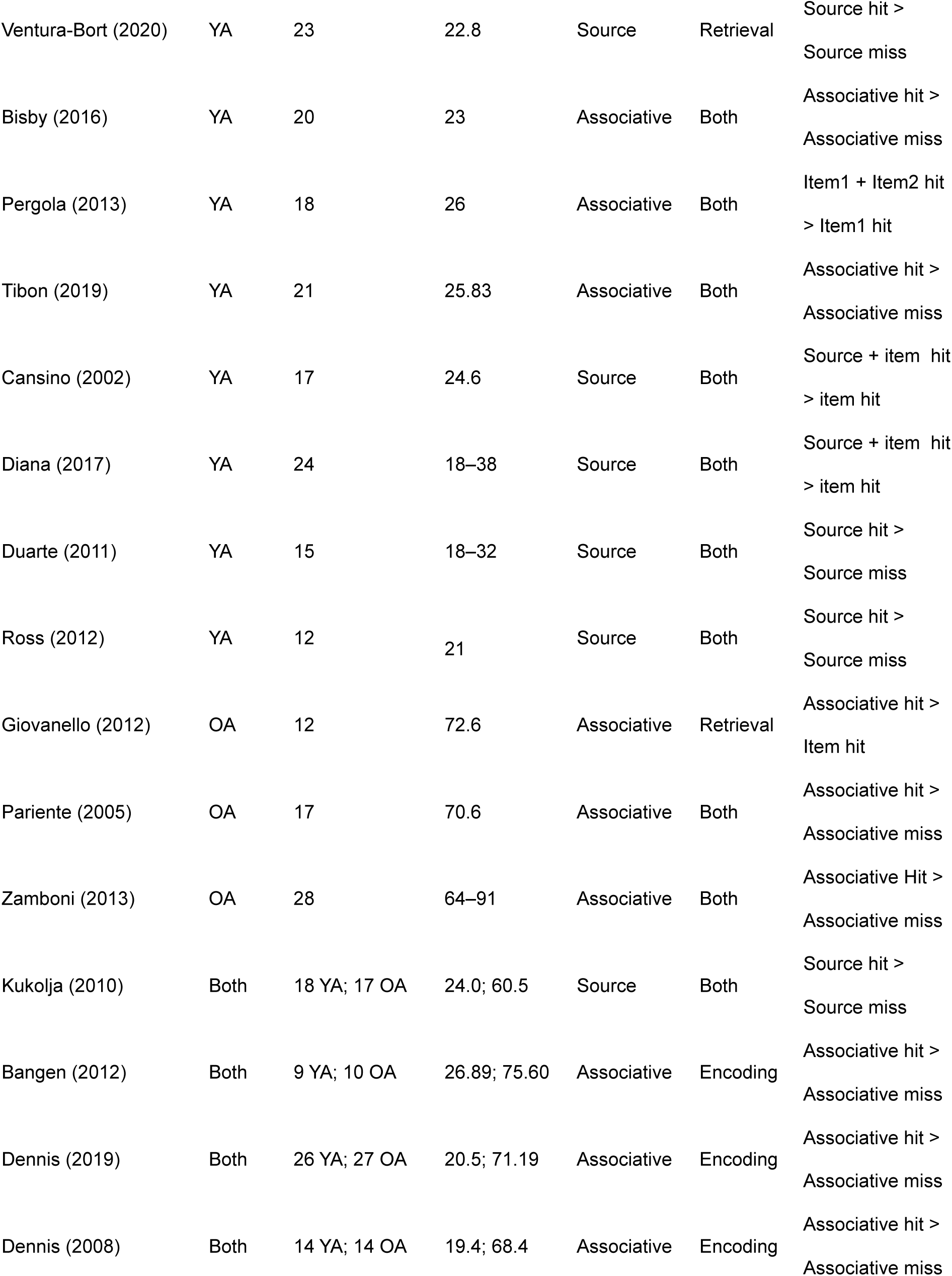
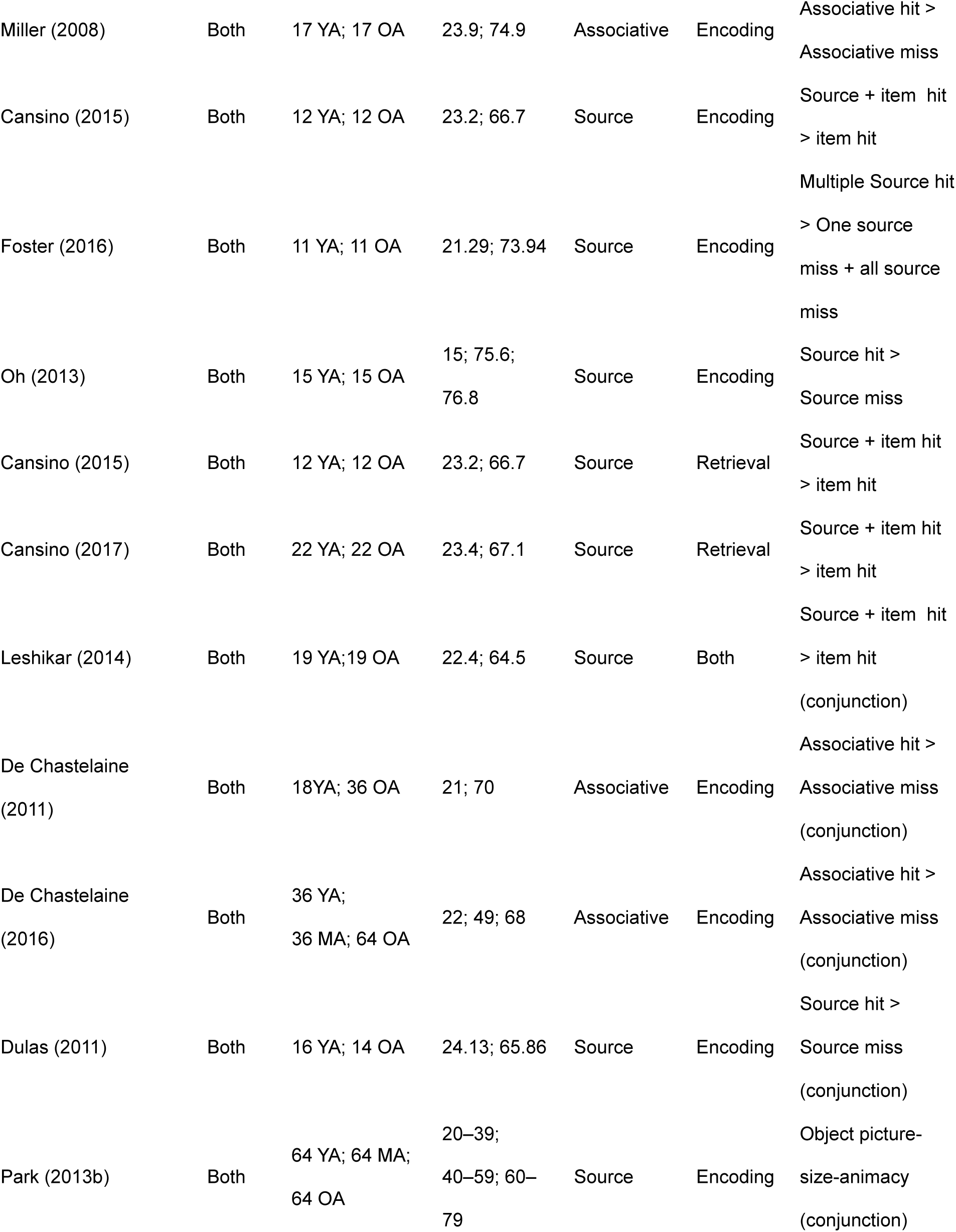
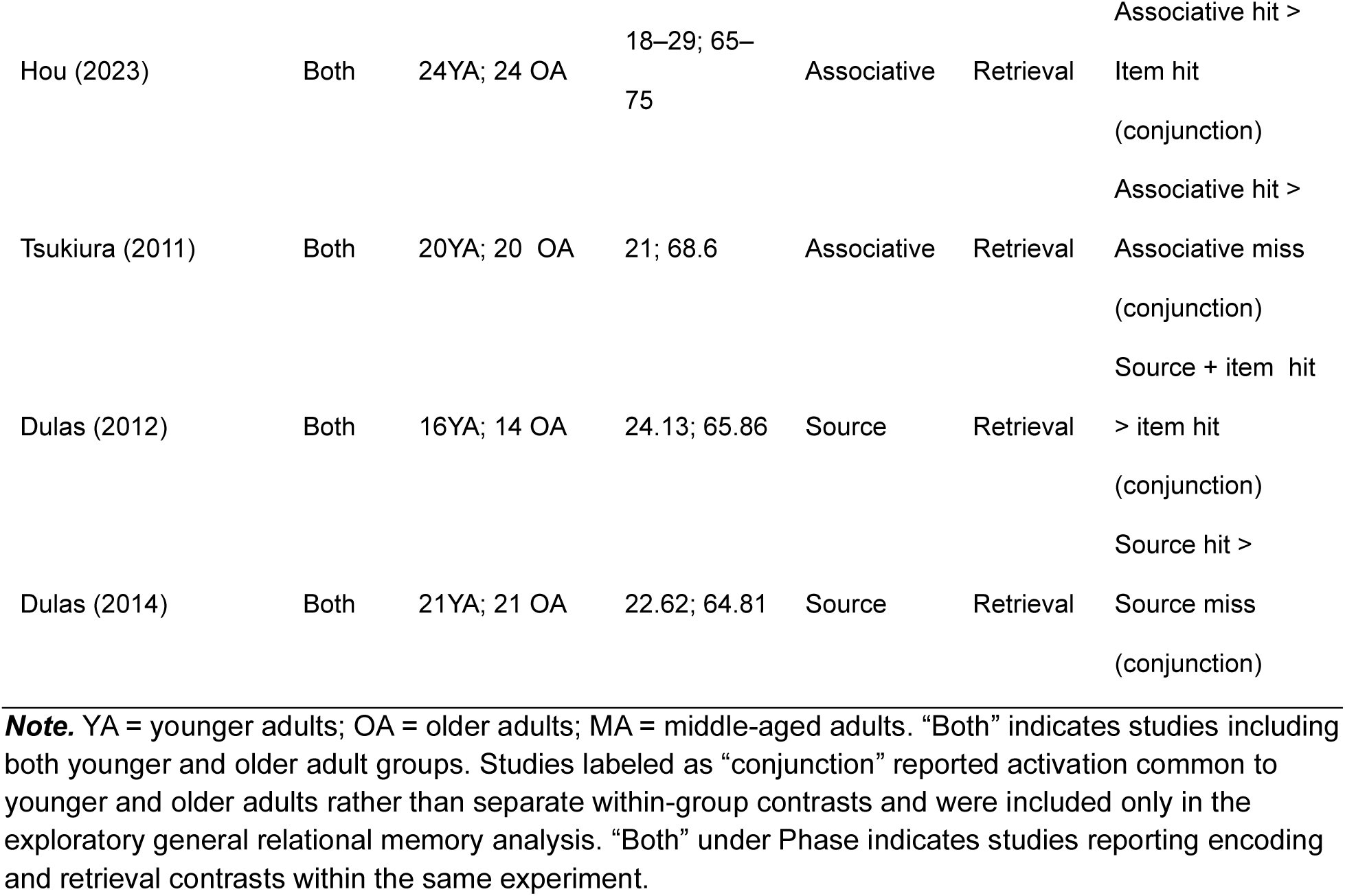
Summary of studies included in the meta-analysis.

**Supplementary list 1.** Separate reference for studies included in meta-analyses

